# PDX Finder: A Portal for Patient-Derived tumor Xenograft Model Discovery

**DOI:** 10.1101/291443

**Authors:** Nathalie Conte, Jeremy Mason, Csaba Halmagyi, Steven B. Neuhauser, Abayomi Mosaku, Dale A. Begley, Debra M. Krupke, Helen Parkinson, Terrence F. Meehan, Carol J. Bult

## Abstract

Patient-derived tumor xenograft (PDX) mouse models are a versatile oncology research platform for studying tumor biology and for testing chemotherapeutic approaches tailored to genomic characteristics of individual patient’s tumors. PDX models are generated and distributed by a diverse group of academic labs, research organizations, multi-institution consortia, and contract research organizations. The distributed nature of PDX repositories and the use of different standards in the associated metadata presents a significant challenge to finding PDX models relevant to specific cancer research questions. The Jackson Laboratory and EMBL-EBI are addressing these challenges by co-developing PDX Finder, a comprehensive open global catalog of PDX models and their associated datasets. Within PDX Finder, model attributes are harmonized and integrated using a previously developed community minimal information standard to support consistent searching across the originating resources. Links to repositories are provided from the PDX Finder search results to facilitate model acquisition and/or collaboration. The PDX Finder resource currently contains information for more than 1900 PDX models of diverse cancers including those from large resources such as the Patient-Derived Models Repository, PDXNet, and EurOPDX. Individuals or organizations that generate and distribute PDXs are invited to increase the “findability” of their models by participating in the PDX Finder initiative at www.pdxfinder.org.

## INTRODUCTION

PDX models recapitulate many of the disease hallmarks of cancer patients and are increasingly being used to study therapeutics, tumor evolution, and drug resistance mechanisms. PDX models are typically generated by the implantation of human tumor tissues or cells into severely immunodeficient mouse hosts. Tumors that engraft successfully are passaged further to generate cohorts of tumor-bearing mice for experimental studies. PDX models are generated and used by researchers in university, clinical, and pharmaceutical industry settings as well as specialized commercial organizations. International consortia focusing on the use of PDX models including PDX-Net and EurOPDX have recently been funded ensuring PDX models will continue to be an important contributor to understanding and treating cancer.

The distributed and diverse nature of PDX repositories as well as differences in associated metadata describing the models presents a significant challenge to researchers seeking to find PDXs that are relevant to specific cancer research questions. To address this issue The Jackson Laboratory and the European Molecular Biological Laboratory-European Bioinformatics Institute (EMBL-EBI) have implemented PDX Finder (http://www.pdxfinder.org), a freely available and searchable catalog of global PDX repositories. The data model for PDX Finder is based on the minimal information standard for PDX models developed in collaboration with a broad range of stakeholders who create and/or use PDX models in basic and pre-clinical cancer research (1). PDX Finder currently provides access to information about more than 1900 PDX models in 7 repositories around the world, including NCI’s Patient Derived Model Repository, The Jackson Laboratory’s PDX Resource, members of the EurOPDX Consortium, and members of NCI’s recently launched PDXNet. Here we describe the implementation of PDX Finder and illustrate how it can be used to locate relevant PDX models. We also provide information on how investigators with small or large repositories of PDX models can have their resource indexed in PDX Finder to improve the visibility of their models.

## PDX FINDER FEATURES AND FUNCTIONALITY

PDX Finder was designed and implemented following a series of workshops and on-line surveys aimed to identify the diverse needs of researchers who use PDX models. The three top-rated functions/interfaces to emerge from the requirements gathering process were:

1. Availability of high level summaries (graphical and tabular) that provide an overview of all the models and repositories included in PDX Finder.
2. Search forms that allow researchers to find PDX models based on cancer type, diagnosis, availability of specific datasets, molecular markers, and results from drug resistance/sensitivity dosing studies.
3. Summary pages for individual models that include salient details about how the model was generated and links to available data sets and model acquisition information.

### PDX Finder home page

The PDX Finder home page (Figure 1) prominently features search capabilities allowing users to perform a lexical search over a number of categories and summary graphics of the resource. The number of PDX models is represented by a list of models by cancer type as well as a pie chart grouping models by anatomical system. A list of all model providers is also featured. The top menu bar provides details about the PDX Finder project, including our information gathering process, contact information, and instructions on how to submit PDX model information to the resource. A “News and Events” section on the bottom of page provides headline summary and links describing news about the PDX resource as well as a Twitter feed where users can follow to be updated on the latest developments.

**Figure 1.**
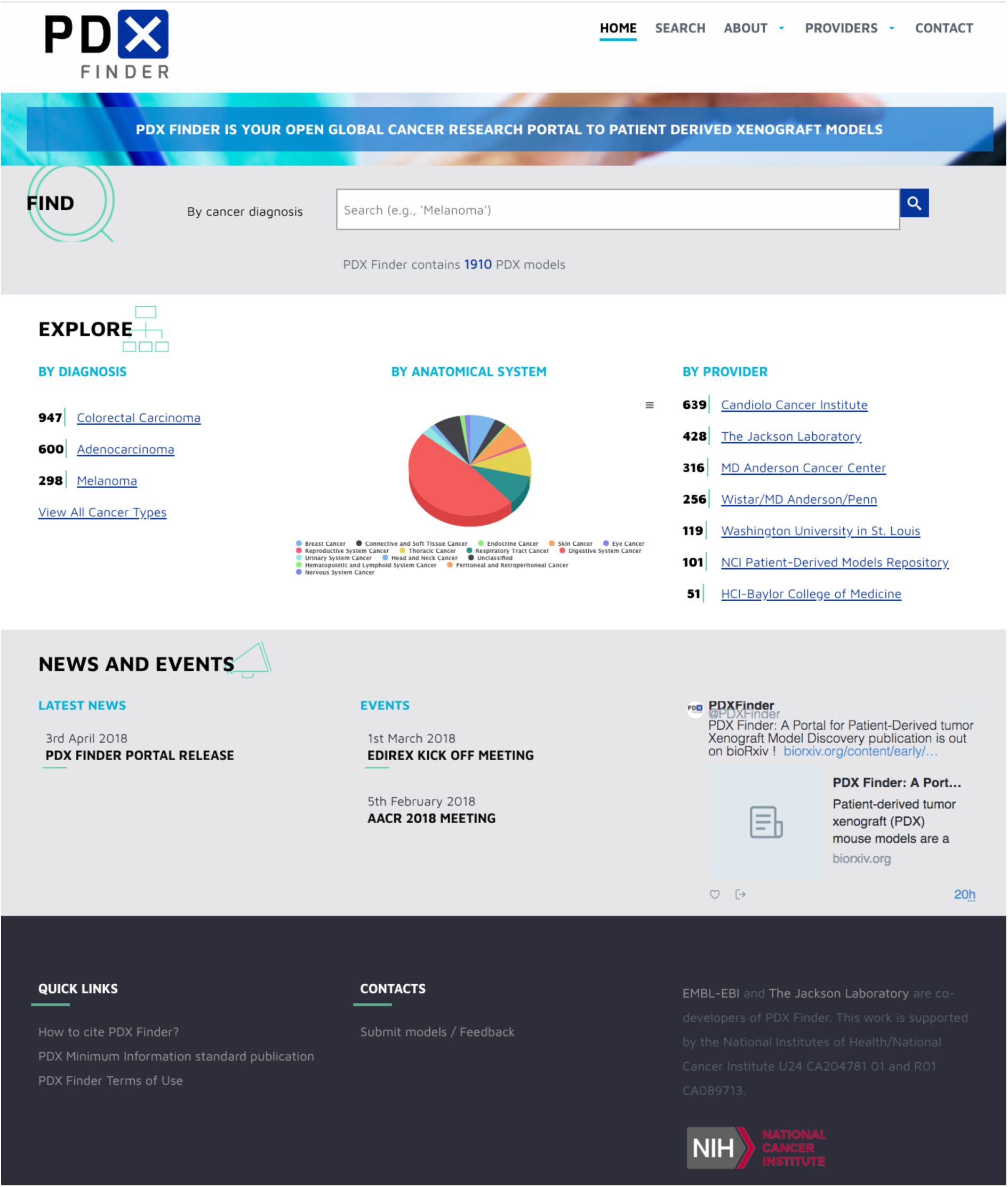
PDX Finder Homepage.

### Search functions

Users navigate to the “Search Results” page by searching for a term or choosing a category from one of the graphical visualizations. Results are presented in a tabular format with each row depicting a model (Figure 2). Key features of the models are presented in columns and include model ID, critical patient clinical data, tumor histological classification, links to available datasets, and the original provider. Multiple filters provided on the left margin of the page allow users to exactly specify the models they would like to appear in the results table. Filters are selected by expanding on a category and selecting the relevant sub-categories with the ability to choose multiple filters. The search functionality allows users to narrow model availability by specific criteria such as “Find all colorectal cancer models with BRAF V600E mutation,” which will retrieve 31 models from 2 different sources. For searches that return multiple models the results can be exported in a tabular format. Users can navigate to a model page or to the data of interest by clicking on the unique PDX Model ID or data links in each row.

**Figure 2.**
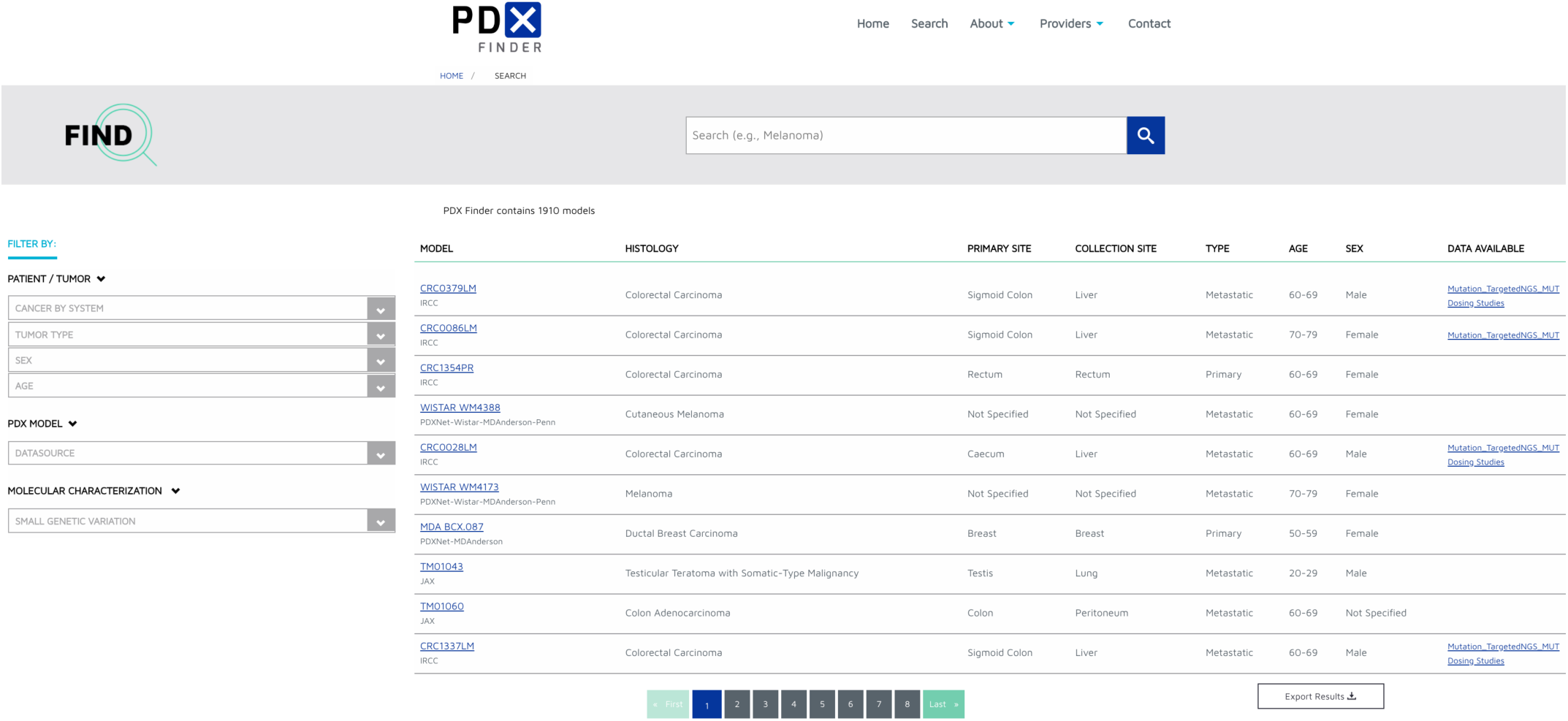
PDX Model Search page.

### Summary pages

The PDX Model Detail page (Figure 3) presents key features about the model. The top of the page displays the model ID and tumor histological classification as well as prominent links to the originating resource where users can find more information and contact the relevant institution for further collaboration. A tabulated section beneath the overview provides summary views of the models with clinical, model, and validation information as well as views of differing types of data submitted with from the PDX model such as genomic or dosing studies when provided by the source.

**Figure 3.**
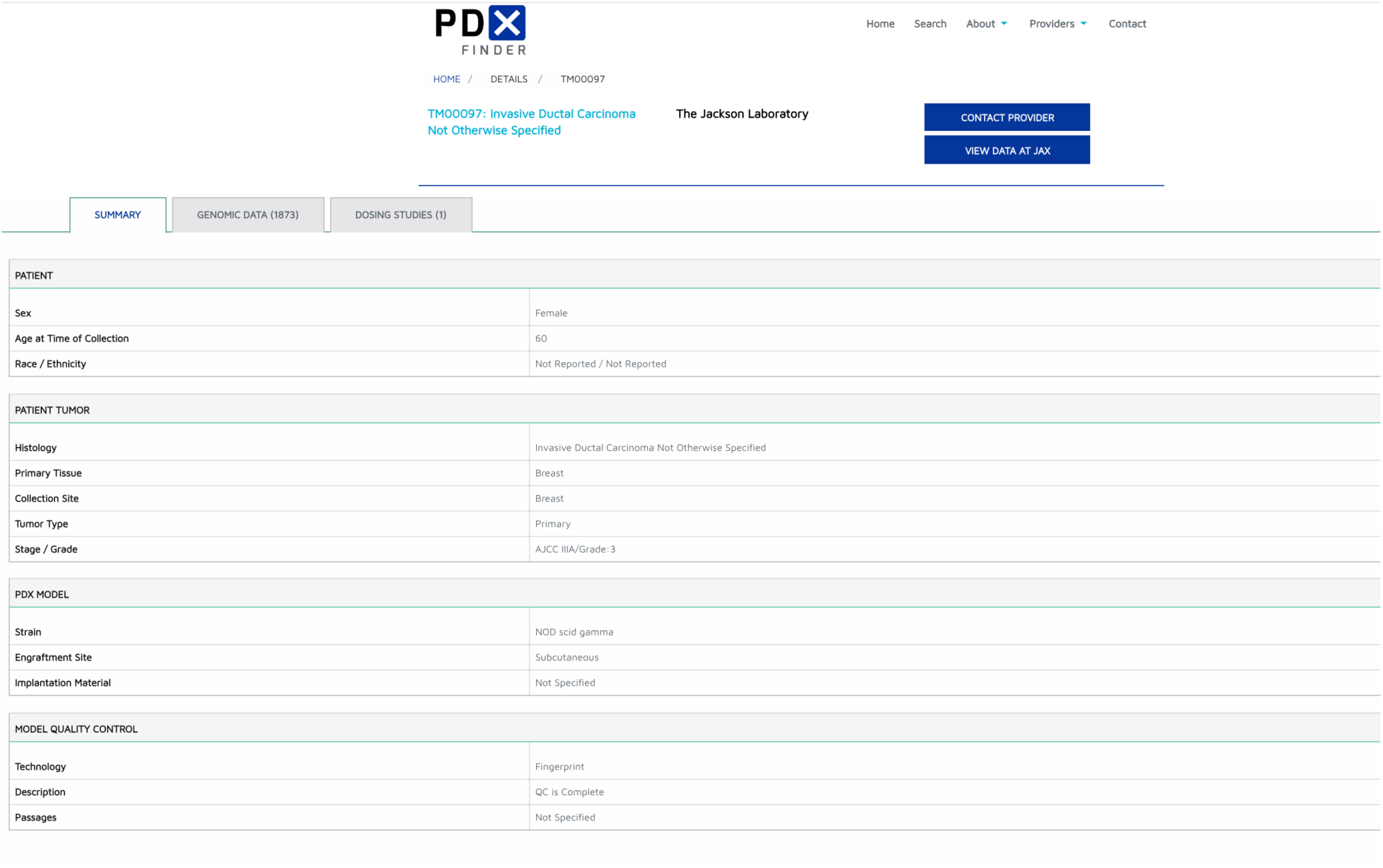
PDX Model Detail page.

## NOMENCLATURE AND METADATA STANDARDS

PDX Finder facilitates discovery of similar models across resources by the use and enforcement of nomenclature and metadata standards. Cancer type, diagnosis, and other cancer attributes are represented by the NCI Thesaurus, a community resource that is actively maintained by the NCI (2). Names and symbols for human genes follow the nomenclature standards approved by the HUGO Gene Nomenclature Committee (3). Host mouse strain nomenclature follows the official guidelines established by the International Committee on Standardized Genetic Nomenclature for Mice (4). Drug/compound names use standards from CHEBI (5), CHEMBL (6), and PubChem (7). In addition, we provide feedback to ontology developers to improve these resources for representing the complexities of PDX models.

## DATABASE AND SOFTWARE INFRASTRUCTURE

The PDX Finder web site was implemented using a combination of the WordPress content management system and a Java web application. The database for PDX Finder is an instance of a Neo4J graph database. To populate this database bespoke Extraction, Transformation, Loading (ETL) software pipelines were written in Java to extract relevant attributes corresponding to the PDX minimal information standard from the data provided by the different PDX repositories. Software developed by the PDX Finder team is freely available under an Apache 2 license (https://www.apache.org/licenses/LICENSE-2.0). The PDX Finder source code is available at GitHub (https://github.com/pdxfinder).

A number of key attributes specified in the minimal information standard rely on different vocabularies and ontologies. To address this challenging harmonization issue PDX Finder employs a semi-automated integration approach using resources at EMBL-EBI including the ZOOMA annotation tool (8) that maps free text annotations to ontology terms based on a curated repositories of annotation knowledge. Transformed metadata is reviewed by local experts and reported to the submitters for validation and approval. Once approved, the transformed metadata describing a model is then loaded into the PDX Finder database and exposed to the users through the web interface.

An example of metadata harmonization includes mapping biologically identical histological concepts harboring different names provided by several sources using their own classification. To harmonize those concepts and support consistent searching across the originating resources, we use different attributes like the original histological term and the primary tissue provided by the source to refine the subtype. Examples of harmonization include mapping source specific histological concepts (“Adenosquamous”, “adenosquamous carcinoma”, “Ad and SC carcinoma”) that share the same primary tissue (“lung”) to the NCIT ontological label “Adenosquamous Lung Carcinoma”. Furthermore, ontological association allows aggregation of concepts based on meaningful groupings like cancer by anatomical system or cell morphology. This allows users to search for “Lung Cancer” models across all subclasses of lung cancer models in a single query, without having to look for each subtype individually.

## FUTURE DIRECTIONS

PDX Finder is a freely available global catalog of PDX models and associated data available from independent, distributed repositories. Future development of PDX Finder will focus on three areas: addition of new PDX repositories, coordination with other informatics groups, and implementation of user-requested functionality.

### Adding new PDX repositories

We will increase the number of PDX models represented in the resource by contacting PDX collection owners. To support scalability of data submission and processing to PDX Finder, we will develop new simplified data submission tools to encourage data submission via spreadsheets or via APIs. The existing ETL modules will be improved by addition of curated knowledge bases that improve the automated data mappings as well as interfaces that facilitate review of the resulting data integration. We will also leverage EBI’s Ontology Xref service (OxO; https://www.ebi.ac.uk/spot/oxo/) that captures community defined cross references between semantic standards in our ETL modules such as mappings between MeSH AND NCI Thesaurus (9).

### Coordination with other informatics resources

To facilitate data sustainability, maximize use of existing data, and avoid redundancy, the PDX Finder team has initiated coordinating activities with existing molecular archives to deposit data generated from PDX Models. The data loading process will have oversight by a PDX Data Coordinator who will broker submission of data to established archives in collaboration with the data providers. Data that have the potential to allow identification of individuals will be submitted to relevant secure molecular archives such as the European Genome-Phenome Archive (10) or dbGAP (11). These resources provide secure environments where the approval of a submitter defined Data Access Committee is required for data access. Depending on the data type, non-patient identifiable data will be submitted to other archive resources including the Sequence Read Archive (12) or the European Nucleotide Archive (13) for nucleotide sequencing information, Gene Expression Omnibus (14) or ArrayExpress (15) for gene expression data, or European Variation Archive (https://www.ebi.ac.uk/eva/) for genetic variation information. PDX Finder will also coordinate the submission of sample metadata to BioSamples database (16), or BioSample archive (17), which provides unique identifiers that are used to link varying types of data derived from the same sample. Integration of the PDX Finder datasets linked to their harmonized data in molecular archives will ensure that meta-analysis type of studies combine biologically comparable datasets from multiple sources.

### New functionality

Other areas of development will be determined by the needs of the PDX research community. We will continue to assess user needs by surveys and testing of the portal. As the number of informatics resources using PDX models grows, the community is best served by sharing solutions to common challenges. The developers of PDX Finder are providing software components to the PDXNet and the EurOPDX data platforms, which is facilitating knowledge exchange and possible reuse of informatics tools. PDX Finder is also funded in part by NCI’s ITCR program, a trans-NCI initiative to promote integration of informatics technology development and reuse of software for oncology researchers. We will be working with developers of other ITCR tools to reuse software developed by others to address common needs.

## COMMUNITY OUTREACH AND USER SUPPORT

The PDX Finder user help desk is available via email and user outreach is supported through Twitter and the PDX Finder listserv. Researchers interested in listing their PDX repository in PDX Finder should contact the team using the helpdesk email provided below. PDX Finder is freely available to any group, organization from academia, or industry that seeks to openly distribute PDX models and associated resources and information.

- Email access:
- Twitter: https://twitter.com/PDXFinder
- PDX Finder Listserve: https://listserver.ebi.ac.uk/mailman/listinfo/pdxfinder-announce

## CITING PDX FINDER

For a general citation of PDX Finder, researchers should cite this article. In addition, the following citation format is suggested when referring to specific data about PDX models obtained from the PDX Finder web site: PDX Finder (http://www.pdxfinder.org); data retrieved in April 2018.

## ACKNOWLEDGEMENTS

The authors thank the PDX resources who have contributed PDX metadata and information to PDX Finder, including members of PDXNet (https://www.pdxnetwork.org/), members of EurOPDX (http://europdx.eu/), The Jackson Laboratory PDX Resource (http://tumor.informatics.jax.org/mtbwi/pdxSearch.do), and NCI’s PDMR (https://pdmr.cancer.gov/). We are particularly grateful to our colleagues, Drs Andrea Bertotti, Enzo Medico (University of Torino, Italy), Aikaterina Chatzipli, Ultan McDermott (Wellcome Sanger Institute, UK), and Yvonne Evrard (NCI Patient-Derived Models Repository, USA) who have provided their expertise and guidance during all stages of the work.

